## Supplementary figures and images for "25 years of propagation in suspension cell culture results in substantial alterations of the *Arabidopsis thaliana* genome"

### Figure S1

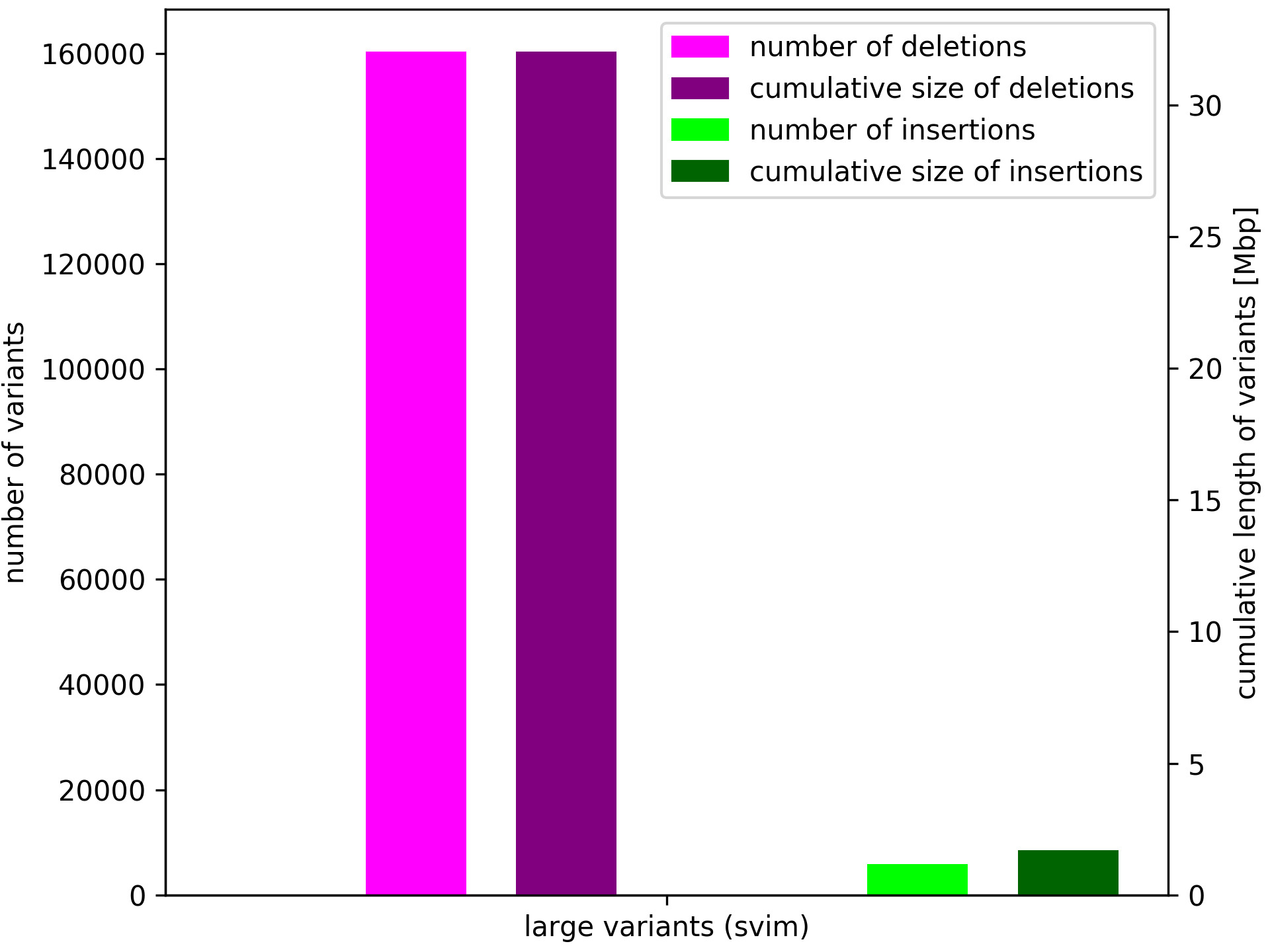

### Figure S2

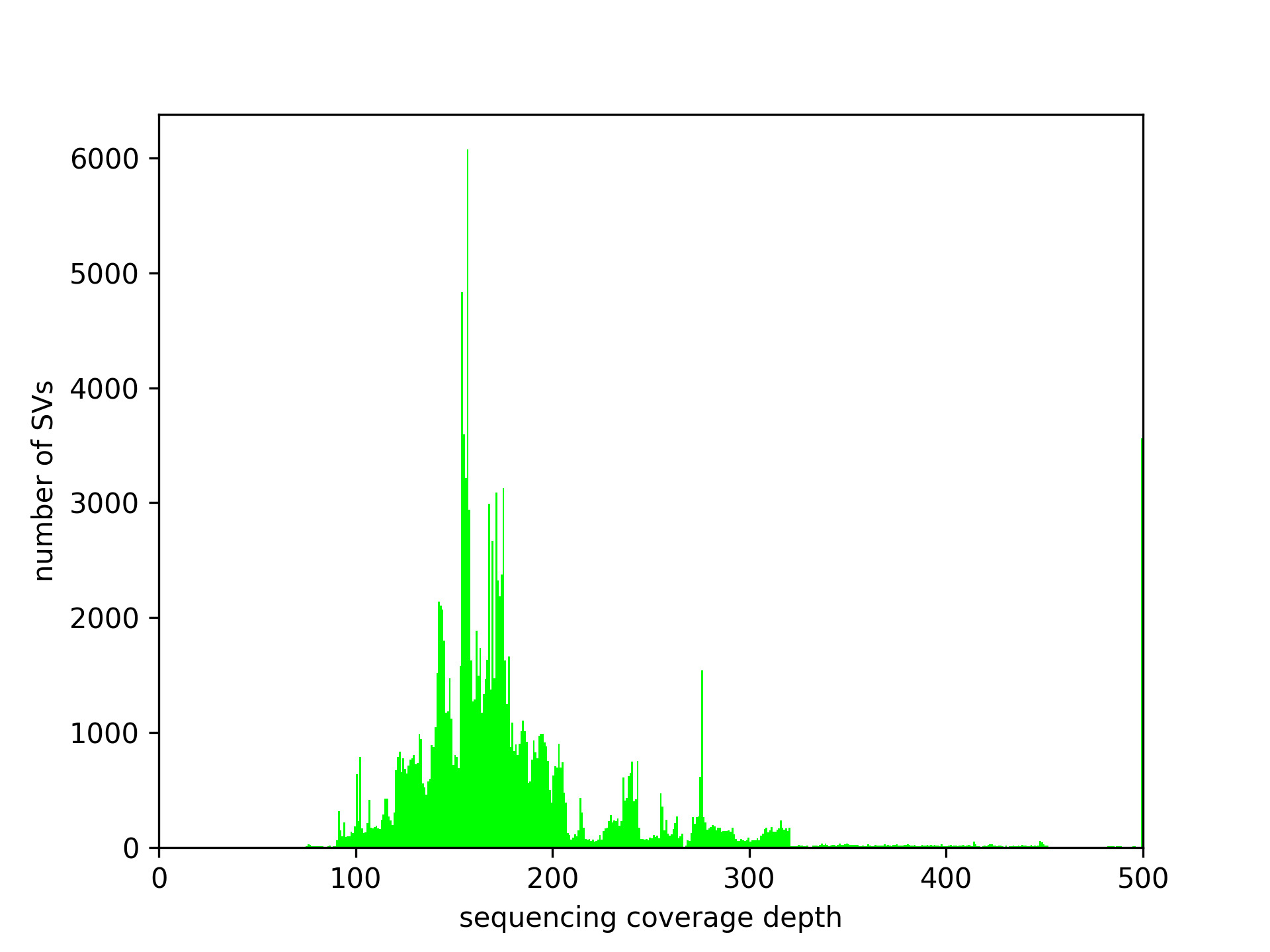

### Figure S3

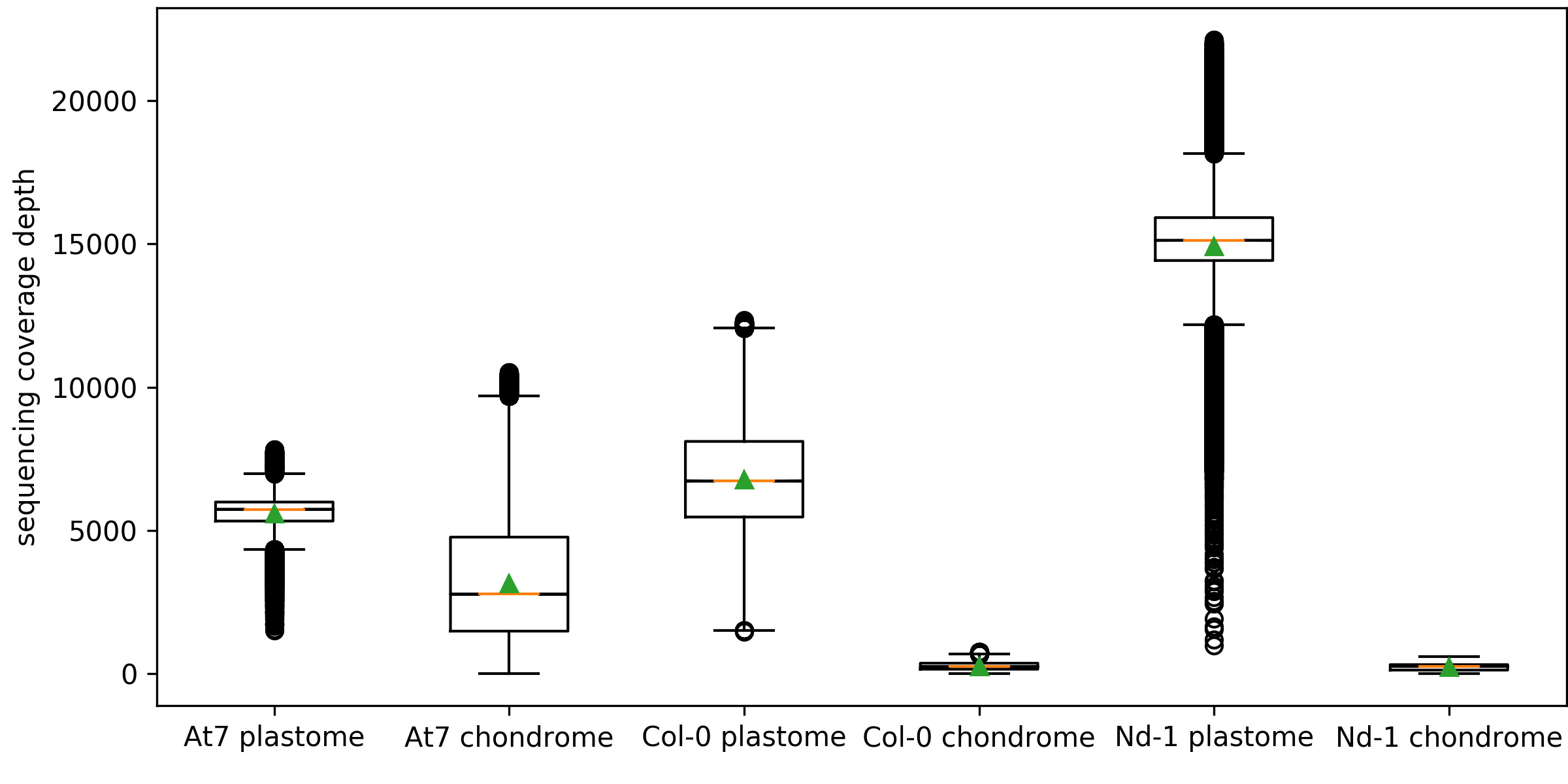

### Figure S4

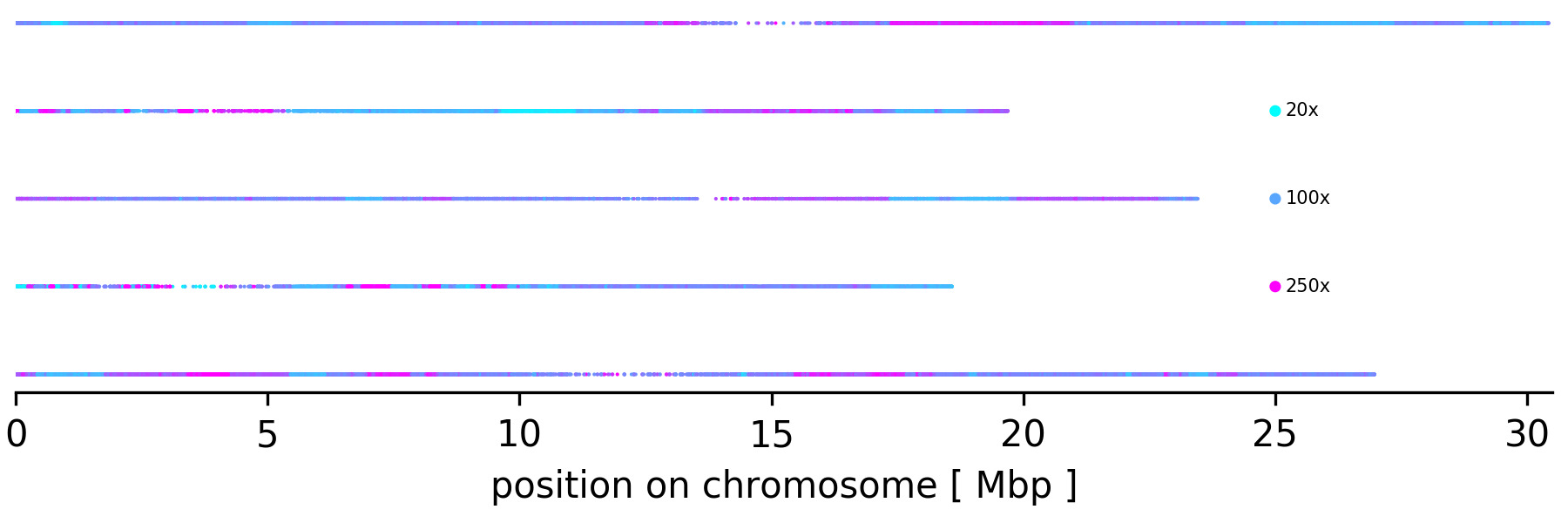

### File S1

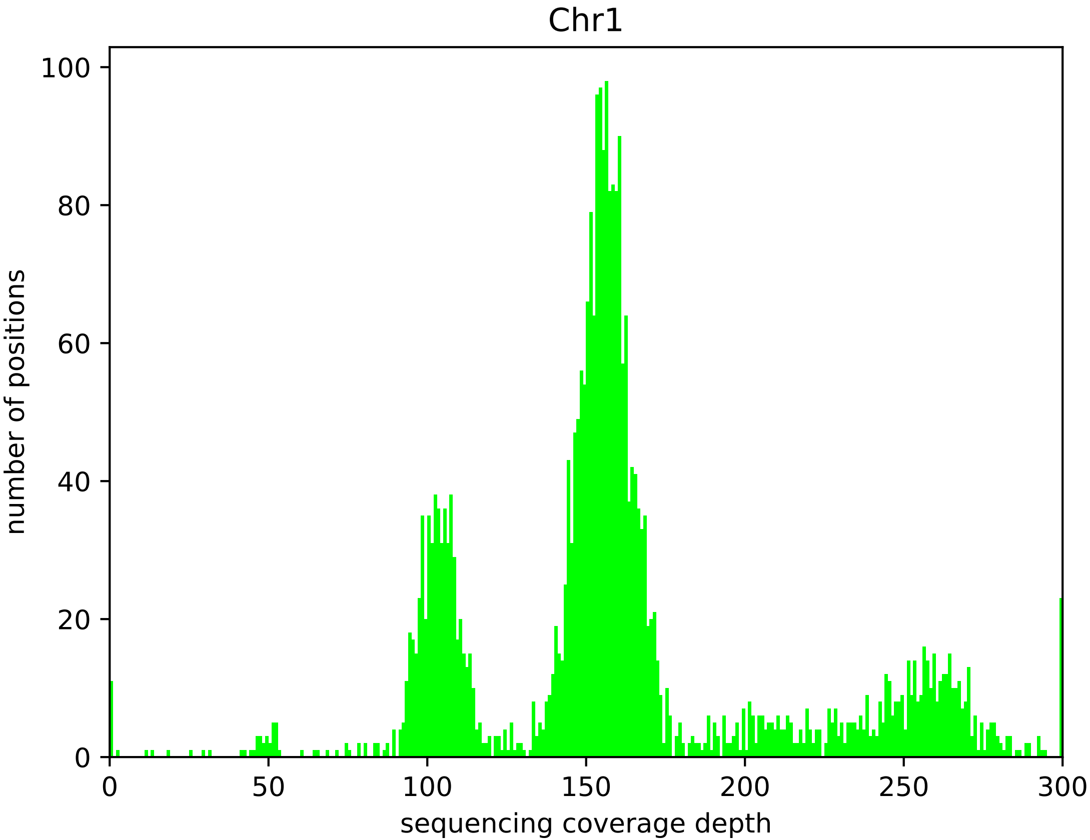

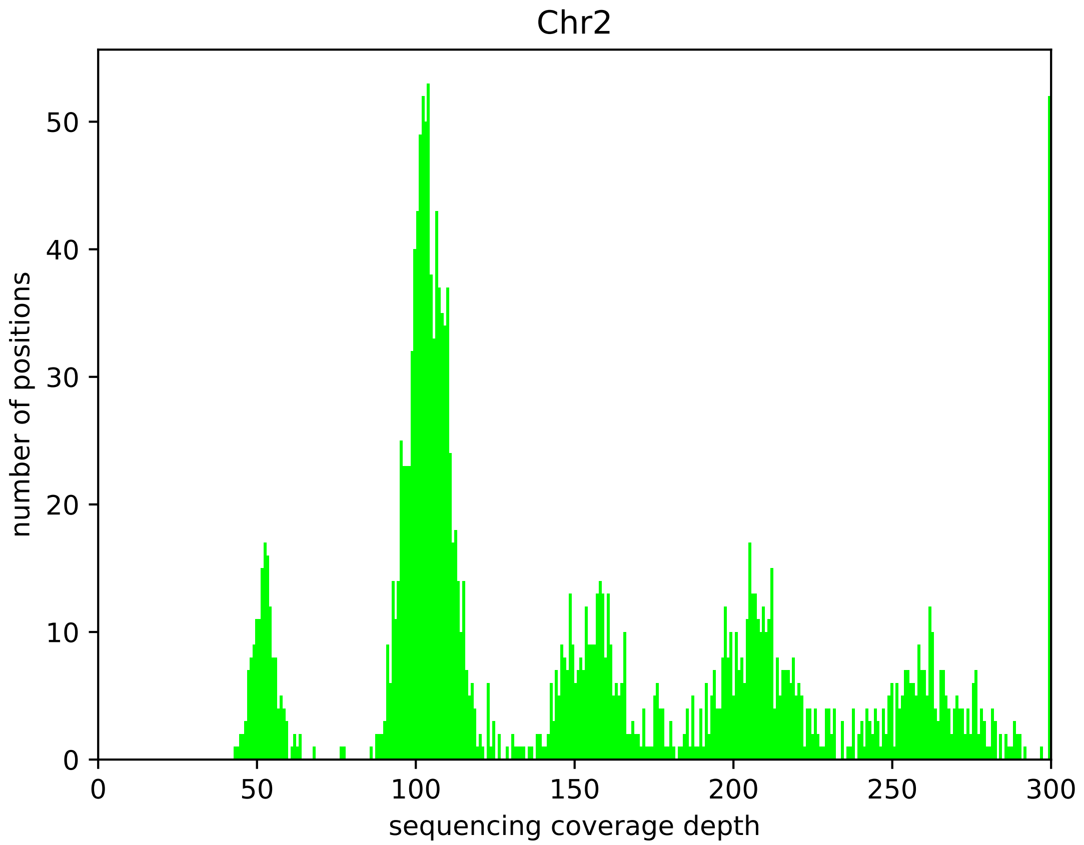

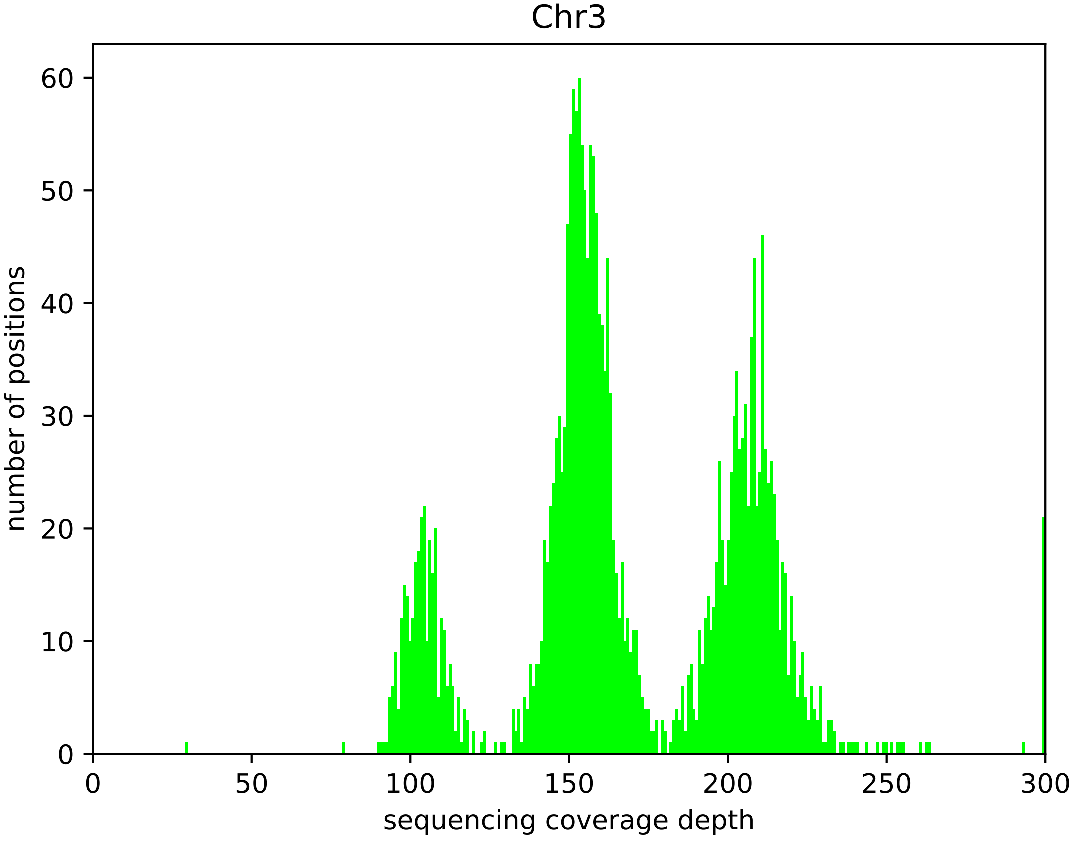

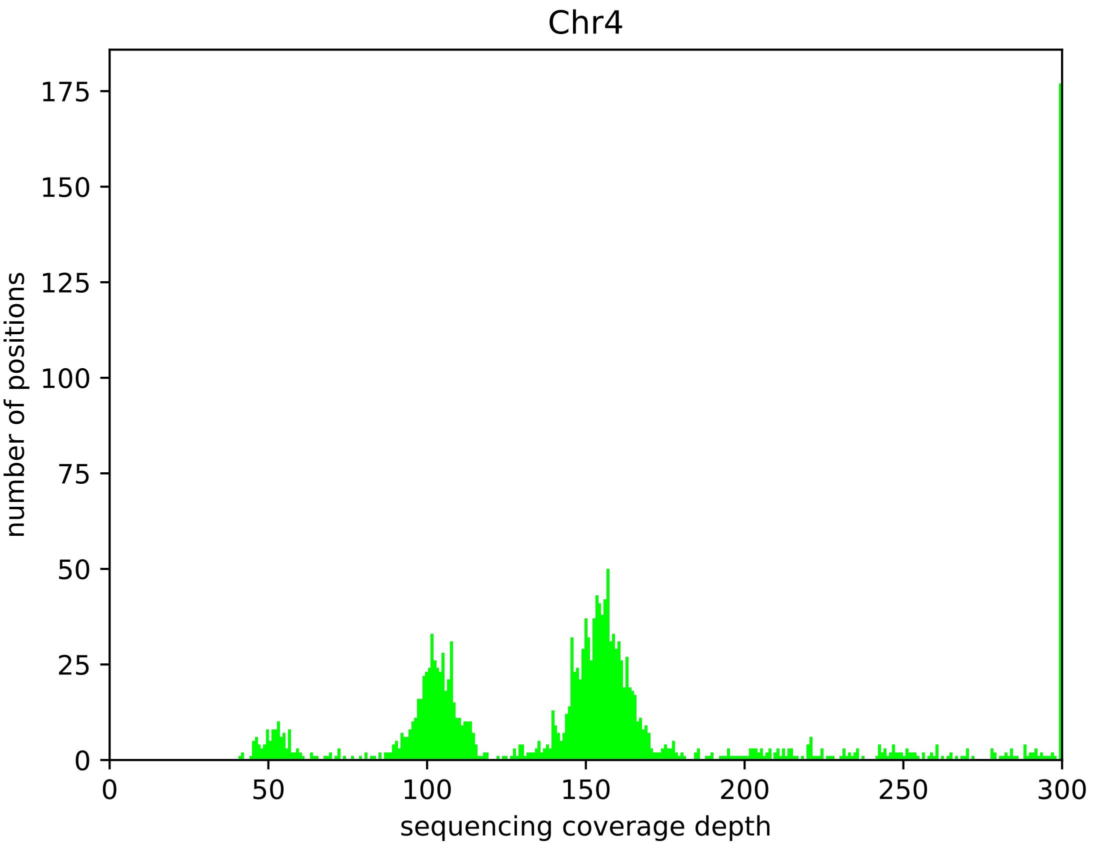

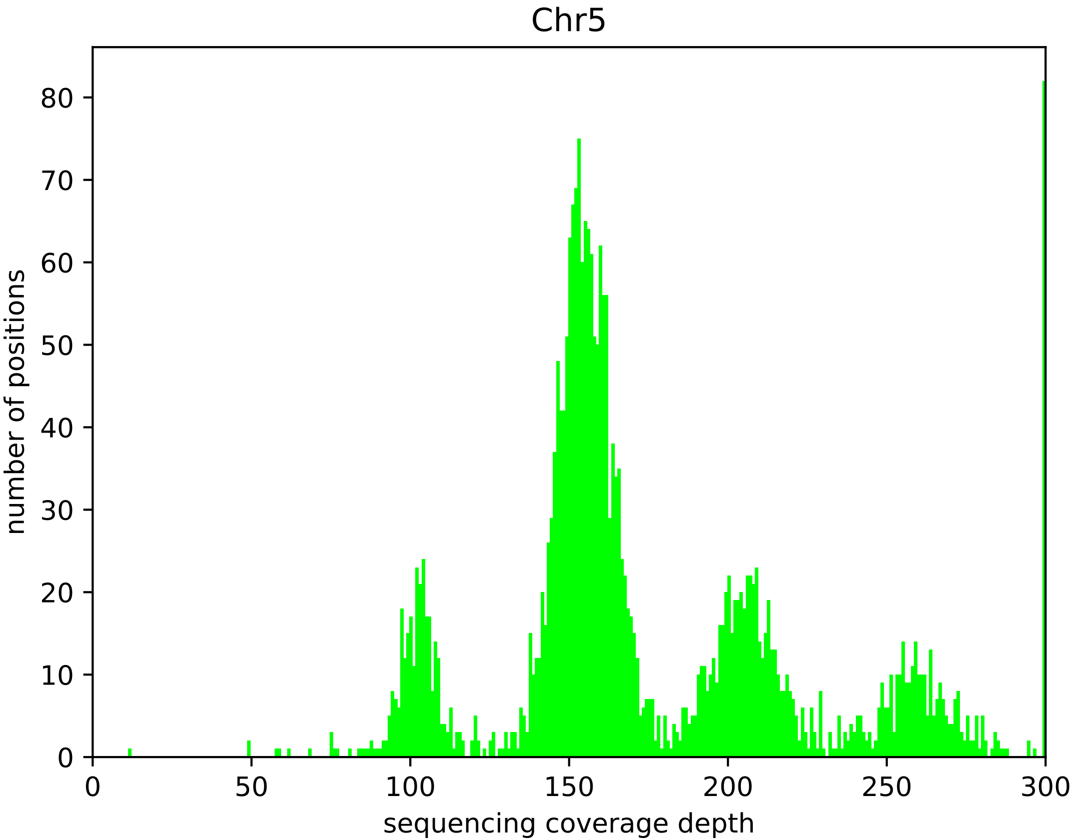
